# Conditions for numerically accurate TMS electric field simulation

**DOI:** 10.1101/505412

**Authors:** Luis J. Gomeza, Moritz Dannhauera, Lari M. Koponena, Angel V. Peterchev

**Author notes:** Corresponding Author: Angel V. Peterchev, Address: 40 Duke Medicine Circle, Box 3620 DUMC, Durham, NC 27710.

## Abstract

**Background:** Computational simulations of the E-field induced by transcranial magnetic stimulation (TMS) are increasingly used to understand its mechanisms and to inform its administration. However, characterization of the accuracy of the simulation methods and the factors that affect it is lacking.

**Objective:** To ensure the accuracy of TMS E-field simulations, we systematically quantify their numerical error and provide guidelines for their setup.

**Method:** We benchmark the accuracy of computational approaches that are commonly used for TMS E-field simulations, including the finite element method (FEM), boundary element method (BEM), finite difference method (FDM), and coil modeling methods.

**Results:** To achieve cortical E-field error levels below 2%, the commonly used FDM and 1^st^ order FEM require meshes with an average edge length below 0.4 mm, whereas BEM and 2^nd^ (or higher) order FEM require edge lengths below 1.5 mm, which is more practical. Coil models employing magnetic and current dipoles require at least 200 and 3,000 dipoles, respectively. For thick solid-conductor coils and frequencies above 3 kHz, winding eddy currents may have to be modeled.

**Conclusion:** BEM, FDM, and FEM methods converge to the same solution. However, FDM and 1^st^ order FEM converge slowly with increasing mesh resolution; therefore, the use of BEM or 2^nd^ (or higher) order FEM is recommended. In some cases, coil eddy currents must be modeled. Both electric current dipole and magnetic dipole models of the coil current can be accurate with sufficiently fine discretization.

**Funding:** Research reported in this publication was supported by the National Institute of Mental Health and the National Institute of Neurological Disorders and Stroke of the National Institutes of Health under Award Numbers RF1MH114268 and R01NS088674-S1. The content of current research is solely the responsibility of the authors and does not necessarily represent the official views of the National Institutes of Health.

**HIGHLIGHTS:**

- FDM and 1^st^ order FEM with 1.5 mm average mesh edge length have numerical errors above 7%.
- BEM or 2^nd^ order FEM are most efficient for achieving numerical errors < 2%.
- Coil wire cross-section must be accounted to achieve E-field errors below < 2%.
- Coil eddy currents can account for > 2% of E-field when very brief pulses are used.

## INTRODUCTION

During transcranial magnetic stimulation (TMS) application, a coil placed on the scalp generates a magnetic field that induces an electric field (E-field) in the underlying head and brain [1]. The delivered E-field dose mediates the physiological effects of TMS; however, there are no established methods for directly measuring the E-field. Therefore, computational modeling has become the dominant approach for TMS E-field dosimetry [2,3].

To ensure safe and effective use of E-field simulations, it is important to systematically quantify their accuracy. TMS simulations require the generation of models of the subject’s head and TMS coil, and their placement relative to each other. The fidelity of the E-field model is typically limited by the accuracy of the tissue conductivity values, the resolution and quality of the head imaging data and its segmentation into discrete tissues, and the accuracy of the coil representation and placement relative to the head [4-9]. These factors can contribute modeling error in the predicted E-field. Additionally, the computational algorithms that calculate the E-field can introduce numerical error. In a correctly implemented simulation, the numerical error should be negligibly small compared to the modeling error. This work aims to provide guidelines for accurate TMS modeling using various computational methods by systematically studying the numerical error of the predicted E-field. Specifically, we study the convergence of predicted cortical E-fields with respect to the following numerical approximation variables: coil current approximation method, type of computational method, head mesh density, and computational method approximation order.

For computational modeling of TMS, the induced E-field can be partitioned into a primary E-field due to the coil currents and a secondary E-field due to charge accumulation on regions where there is a change in conductivity [10]. Numerical error of the primary E-field is dependent on the method used for modeling the TMS coil windings and its currents. Numerical error of the secondary E-field is dependent on the numerical method used—finite element method (FEM) or boundary element method (BEM)—and the specific type of FEM or BEM discretization.

Typical TMS coils consist of circular windings of wire with rectangular cross-section. For example, the standard Magstim 70mm Double coil consists of a pair of 70 mm diameter, nine-turn circular spiral windings of wire with cross-section of 1.9 mm by 6 mm. Because of their small cross-section most TMS modeling frameworks assume that the currents flow uniformly through the wire [11-13]. However, the eddy currents cause the currents to redistribute and concentrate toward the wire surface [14]. Neglecting this eddy-current redistribution can potentially result in errors of the predicted primary E-field. Furthermore, some modeling frameworks use Faraday’s law to model the coil as magnetic dipoles, thereby, introducing another possible source of numerical error [11,15,16]. We evaluate different approximations of modeling the coil primary E-field, including the impact of eddy currents and explicit representation of the coil current flow in contrast to approximation through magnetic dipoles.

In FEM, the head is discretized into a mesh consisting of polyhedral volumetric elements [17] (e.g., tetrahedrons or hexahedrons). Then, a weak form of Poisson’s equation [11] is solved numerically to determine the secondary E-field. A commonly used subset of FEM called finite difference method (FDM) approximates the head by a mesh consisting of equal sized cuboid elements, thereby approximating tissue boundaries by ‘staircases’ [18,19]. These staircase approximations can be problematic since accurately resolving tissue boundaries requires many elements [19], and artifactual outliers are typically observed in the FDM solution [18-21]. In contrast, tetrahedral meshes can be used to better conform to the smooth tissue boundaries, and many existing software packages use them for TMS modeling [11,22-24].

For BEM, the subject’s head tissue boundary surfaces are discretized by two-dimensional polyhedral elements (triangles or rectangles). Weak forms of surface integral equations are solved to determine the TMS induced E-field. FEM and BEM solve a system of equations for a piecewise polynomial approximation of a quantity related to the secondary E-field (e.g., charge distributions or scalar potential) on the mesh. Because BEM only discretizes tissue surfaces, it typically requires less elements than FEM. Nevertheless, in BEM computer memory requirements increase quadratically with the number of mesh elements, limiting its use [25]. Recently, an implementation of the adjoint double layer BEM [26,27] using fast-multipole-method (FMM) accelerators, which reduces memory requirements to linearly increasing with the number of mesh elements [28], has been shown to be computationally tractable for analyzing realistic TMS scenarios [29]. A remaining limitation of BEM is that, unlike FEM or FDM, it is not effective at modeling highly heterogeneous media such as anisotropic tissue [30]. For highly heterogeneous media, extensions of BEM termed volume integral equation methods can be used [24,31,32]. However, like FEM and FDM, these methods need volumetric meshes, which makes them more computationally expensive than BEM [30].

Both FEM and BEM are known to converge to the actual solution by either refining the mesh (H-refinement) and/or increasing the polynomial approximation order (P-refinement) [33]. Because of the relatively complex geometry of the human brain, it is difficult to estimate the P- and H-refinement necessary to compute accurate E-field representations in the cortex with FEM and BEM. Therefore, we also evaluate via simulations the impact of P- and H-refinements on the TMS E-field modeling accuracy with FDM, FEM, and BEM, and compare them to a reference solution.

In summary, in this paper we benchmark various TMS E-field simulation methods, and determine the conditions for achieving specific error levels. We compare the absolute accuracy and computation time of FDM, FEM, and BEM implementations with respect to analytically-derived solutions of a spherical head model and a high-fidelity simulation of a realistic head model. We also explore the impact of the TMS coil representation on the solution accuracy.

## METHODS AND MATERIALS

### Approximation of Electromagnetic Equations

We used the quasistatic FEM used by [11,22,23] and FDM used in [18]. In this formulation, displacement currents are neglected, as they are much smaller than conduction currents. Magnetic fields scattered by the human head are also neglected, as they are small compared to the magnetic field produced by the TMS coil. Within these assumptions, the primary E-field can be determined from the coil currents via the Biot-Savart law [34], and the secondary E-field arises purely from a scalar potential field. The scalar potential can be determined by solving Poisson’s differential equation from the primary E-field,

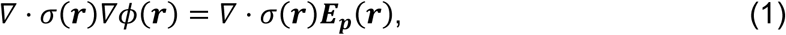

where σ(**r**), ϕ(**r**), and **E**_p_(**r**) are the conductivity, scalar potential, and primary E-field at position **r**, respectively. Both FEM and FDM solve Eq. 1 by approximating the head by a mesh consisting of elements (tetrahedrons for FEM and identical cuboidal elements for FDM), assuming a piecewise polynomial approximation to the scalar potential within each element, and determining the correct piecewise polynomial expansion by applying weak forms of Eq. 1. The convergence of the FEM and FDM approximation with respect to the true solution depends on the size of the individual elements used to approximate the head and the polynomial order of the solution [33]. TMS modeling packages [11,22,23] typically employ piecewise linear (1^st^ order) approximations for the scalar potential, resulting in piecewise constant approximations of the E-field within the head. Relative to higher order methods, these piecewise constant approximations converge slowly to the true solution [33]. Therefore, we additionally evaluated 2^nd^ and 3^rd^ order FEM formulations that result respectively in piecewise linear and piecewise quadratic approximations of the E-field within the head.

For BEM, we used the recently developed FMM-accelerated BEM formulation for analysis of TMS which solves integral equations in terms of surface charges on the tissue boundaries [29]. Then, the surface-charge induced E-field is evaluated using Coulomb's law. We implemented the formulation with a piecewise constant (0^th^ order) [29] as well as a piecewise linear (1^st^ order) approximation of the charges.

All solvers where developed in-house, with implementation details provided in the supplement.

### Head modeling

Spherical head models: we employed concentric spherical shell models [35,36] composed of a spherical core and three spherical shell compartments with inner to outer layers representing cortex, cerebrospinal fluid (CSF), skull, and scalp, with each with respective outer boundary radius of 78, 80, 86, and 92 mm. We considered an inhomogeneous sphere model where the cortex, CSF, skull and scalp layer conductivity is set to 0.33, 1.79, 0.01, and 0.43 S/m, respectively [8]. Additionally, we considered a homogenous sphere model comprised of a single-shell sphere with radius of 92 mm and conductivity of 1 S/m. For both versions of sphere models we used the same FEM meshes, and utilize the meshing software, GMSH [37], for discretization. The resulting mesh parameters are given in Table I. Additionally, we generated FDM meshes centered about the spherical head by assigning each voxel the spherical model conductivity corresponding to the voxel center. For each mesh, we ran a simulation scenario where the head is centered about the origin, and the bottom of the coil windings is centered 5 mm above its apex at location (97, 0, 0) mm, as shown in Fig. 1A. Analytical expressions for the E-field generated inside the spherical head model [38] were used to determine accuracy of all solvers for the sphere head at each mesh density level.

**Figure 1.**
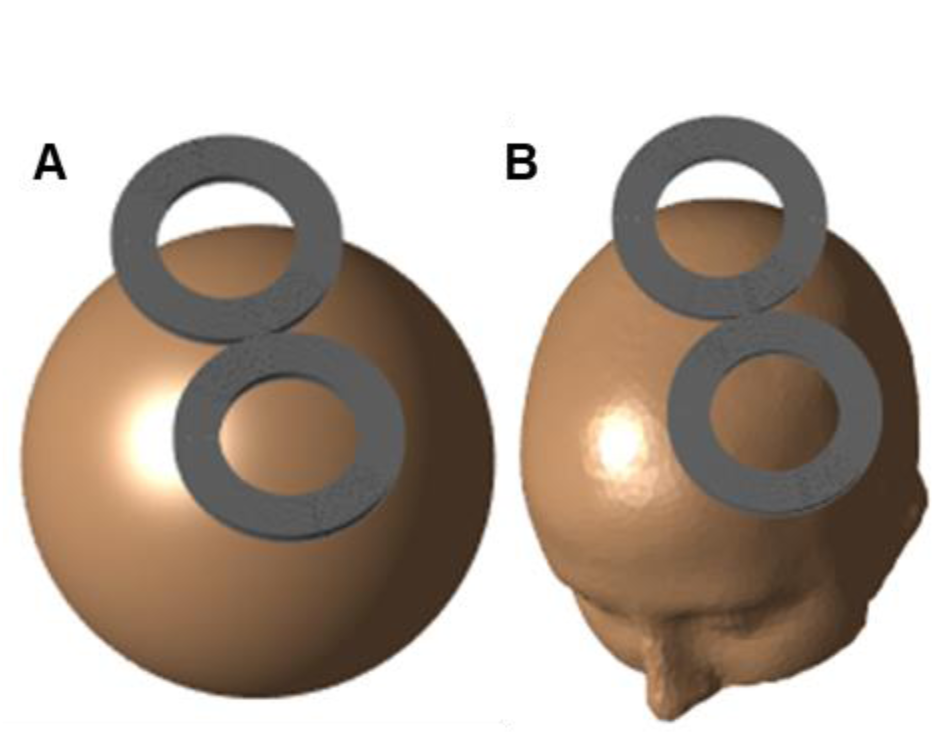
TMS E-field simulation setups. (A) The coil was placed directly above the apex of the spherical head model. (B) The coil was placed above the primary motor cortex hand-knob representation of the MRI-derived head model.

**Table I.** Spherical head model mesh parameters.

| Mesh refinement level | Number of tetrahedrons | Edge length (mm) |  | Number of homogeneous sphere BEM triangles | Number of inhomogenous sphere BEM triangles |
| --- | --- | --- | --- | --- | --- |
|  |  | Average | Maximum |  |  |
| 0 | 11,496 | 14.6 | 32.4 | 1,202 | 3,972 |
| 1 | 54,258 | 8.5 | 17.5 | 4,668 | 15,642 |
| 2 | 319,304 | 4.8 | 9.0 | 20,758 | 68,220 |
| 3 | 1,613,233 | 2.8 | 5.1 | 62,706 | 210,714 |

MRI-derived head model: As a head mesh, we used the one provided in the popular SimNIBS v1.0 package [11]. (In SimNIBS v2.0 this mesh was updated with one having nearly doubled resolution; however, we employed the lower-resolution version since it enables more barycentric refinements while maintaining computational tractability.) The mesh consisted of five homogenous compartments including white/gray matter, CSF, skull, and skin with conductivity of 0.126, 0.276, 1.65, 0.01 and 0.465 S/m, respectively [39]. Starting from the SimNIBS v1.0 example mesh, we generated refined meshes, each time by subdividing each tetrahedron of the earlier mesh into eight equal sized tetrahedrons using the GMSH ‘refine by splitting’ option [37]. Mesh parameters for each refinement level are given in Table II. The coil was placed 5 mm above the primary motor cortex hand-knob region of the brain as shown in Fig. 1B.

**Table II.** MRI-derived head model mesh parameters.

| Mesh refinement level | Number of tetrahedrons | Edge length (mm) |  |  | Number of BEM triangles |
| --- | --- | --- | --- | --- | --- |
|  |  | Minimum | Average | Maximum |  |
| 0 | 877,106 | 0.2 | 2.9 | 15.7 | 251,888 |
| 1 | 7,016,848 | 0.1 | 1.5 | 14.1 | 1,007,552 |
| 2 | 56,134,784 | 0.06 | 0.8 | 7.2 | 4,030,208 |
| 3 | 449,078,272 | 0.03 | 0.4 | 5.3 | 16,120,832 |

### Coil modeling

We considered two models of 70 mm loop diameter figure-8 coils. Both models assume that the coil is made of rectangular cross-section copper wire and consist of two circular windings side by side, where each winding has 9 concentric turns and inner/outer diameter of 52/88 mm, respectively. One model has a ‘thick’ cross-section of 1.9 mm by 6 mm, as measured from a Magstim coil winding. The other model has a ‘thin’ cross-section of 1 mm by 7 mm [16]. Models of the coils were constructed and meshed in GMSH. Same as for the head models, we constructed a hierarchy of meshes each time by using the ‘refine by splitting’ option in GMSH.

The currents flowing through the coil windings were determined by solving the current continuity equation via FEM. Additionally, eddy-current effects were modeled using a time-harmonic partial-element equivalent circuit method [14] developed in-house (details provided as supplementary material). We computed the primary E-field generated by the coil inside the spherical head region using a first order accurate quadrature for each tetrahedron of the coil mesh. Primary E-field samples were taken on a grid with points 2 mm apart. Furthermore, we compared the total E-field predicted by the 2^nd^ order FEM with the MRI-derived head mesh with one refinement level.

Additionally, we modeled the thick cross-section coil with equivalent magnetic dipoles [43]. To generate sets of dipoles with varying density, where each dipole represents a small subregion of the coil, we used dipole-placement rules generalized from the layout used in [19]. First, both circular loops of the figure-8 coil were divided into *N_ring_* rings of uniform width. Then, to obtain most uniform set of dipoles, each ring was discretized with 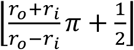. equispaced dipoles (where *r_i_* is the inner radius of the ring and *r_o_*—its outer radius). Lastly, *N_layer_* identical copies of the dipole rings were stacked uniformly between the bottom and top of the coil windings.

For the results comparing head E-field modeling methods, we did not include eddy-effects and computed the primary E-field of the figure-8 coil using an integration rule with 193,536 sample points, which resulted in L^2^ norm error below 0.1% for the primary E-field generated in the head. We used the same coil model for these simulations to remove coil modeling as a contributor to numerical error when comparing E-field solvers.

### Accuracy benchmarks

To quantify the accuracy of our solution we considered two metrics. The first metric is relative L2-norm error,

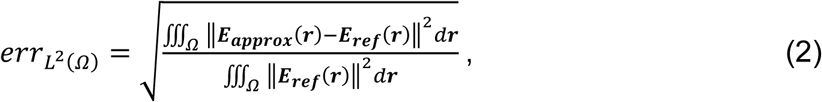

where ‖⋅‖ denotes magnitude, **E**_ref_(**r**) and **E**_approx_(**r**) are the reference and approximate solution, respectively, and Ω is the domain of integration. For head simulations the integration was done over the GM/WM matter region (i.e., Ω = *Brain*) and for coil current error—in the coil winding wire (i.e., Ω = *Wire*). The L^2^ norm error is a measure of the relative energy of the error of the E-field (i.e., the residue), thereby quantifying the global convergence of the numerical solution. For the spherical head model, the cortical L^2^ norm error was estimated by sampling the E-field on a grid with points 2 mm apart. For the MRI-derived head, the cortical L^2^ norm error was estimated by applying a 2^nd^ order accurate integration rule [40] in each tetrahedron of the original mesh.

The second metric is the pointwise magnitude error

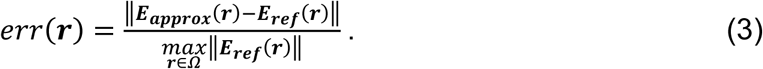

This error quantifies the magnitude of the E-field difference at each point relative to the E-field maximum, thereby, characterizing the local convergence of the numerical solution.

## RESULTS

### Spherical head model

For the inhomogeneous and homogenous sphere simulations, the maximum pointwise E-field error on a spherical shell 0.5 mm into the cortex (i.e., with radius 77.5 mm) and ‘cortical’ L^2^ errors are shown in Fig. 2. For the homogenous sphere, the maximum pointwise and L^2^ errors are similar. FDM, 0^th^ order BEM, and 1^st^ order BEM achieve similar accuracy to 1^st^ order FEM, 2^nd^ order FEM, and 3^rd^ order FEM, respectively. Furthermore, increasing the order of FEM or BEM decreases the L^2^ error by 1 to 2 orders of magnitude.

**Figure 2.**
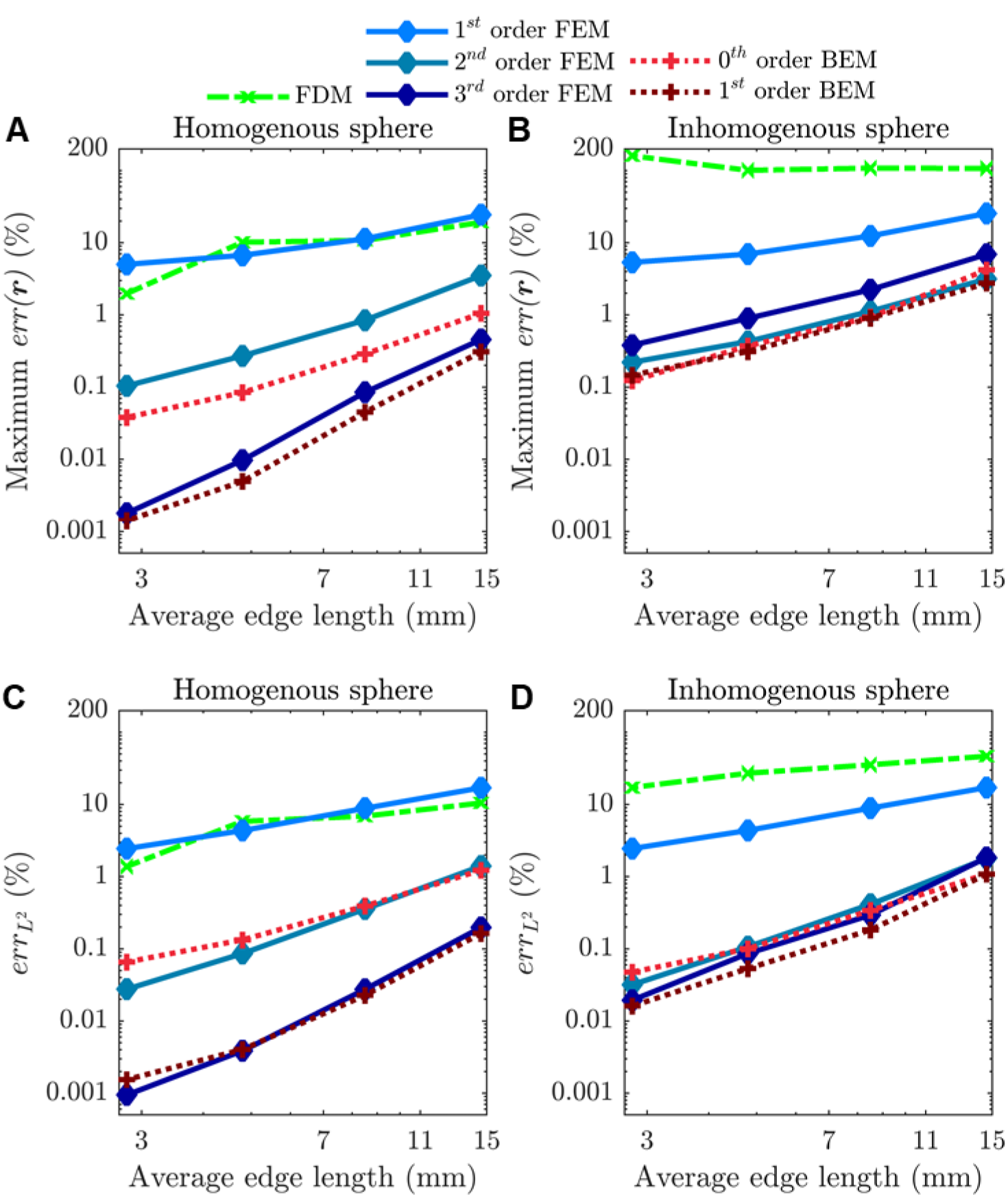
Maximum pointwise error of ‘cortical’ E-field in (A) homogenous and (B) inhomogeneous spherical head model. Global L2 norm error of ‘cortical’ E-field in (C) homogenous and (D) inhomogeneous spherical head model.

Compared to the homogenous sphere (Figs. 2A–C), for the inhomogeneous sphere (Figs. 2B–D) FDM errors increase by at least a factor of 5; 1^st^ order FEM remains virtually unchanged; 2^nd^ order FEM and 0^th^ order BEM errors increase by at most a factor of 2; and accuracy gains from increasing FEM and BEM order from 2^nd^ to 3^rd^ and 0^th^ to 1^st^, respectively, disappear. For FDM a systematic error due to the staircase approximation of each compartment boundary causes the decreased accuracy in the inhomogeneous case and a relatively constant maximum pointwise error with increasing mesh resolution. The low error differences in the inhomogenous case indicate that for orders beyond 2^nd^ for FEM and 0^th^ for BEM, the numerical error near tissue interfaces is dominated by spatial discrepancies between the mesh boundary and the ideal sphere geometry. To achieve significant accuracy gains by increasing order beyond 2^nd^ for FEM or 0^th^ for BEM, it is likely necessary to use higher-order mesh element shape functions [41] to approximate the head geometry.

The sphere is a smooth geometry with minor geometrical features, resulting in a smooth E-field that rapidly converges to the true solution. As a result, we observe errors below 2% even for mesh edge lengths as high as 1.4 cm. The next section analyzes solution convergence in a more realistic head model.

### MRI-derived head model

We consider the E-field simulations for the MRI-derived head models using 0^th^ order BEM and 3^rd^ refinement mesh as a reference solution, since it was the most accurate test case we could run within our computational resources. The pointwise E-field error 0.5 mm into the cortical surface is shown in Fig. 3. For all three refinements, the FDM pointwise error exhibits large regions with errors above 2%; as such, this solution is likely not reliable on this surface. The 1^st^ order FEM solution slowly improves as the mesh is refined. After two refinements, significant regions near the sulci have a pointwise error above 2%, consistent with prior observations [42]. After three refinements, there no longer are significant portions with error above 2%. The three refinements required 400 million elements, in contrast to the typical 3–5 million elements used. All other methods exhibited errors below 2% after one refinement, and after two refinements most of the cortical shell had errors below 0.1%.

**Figure 3.**
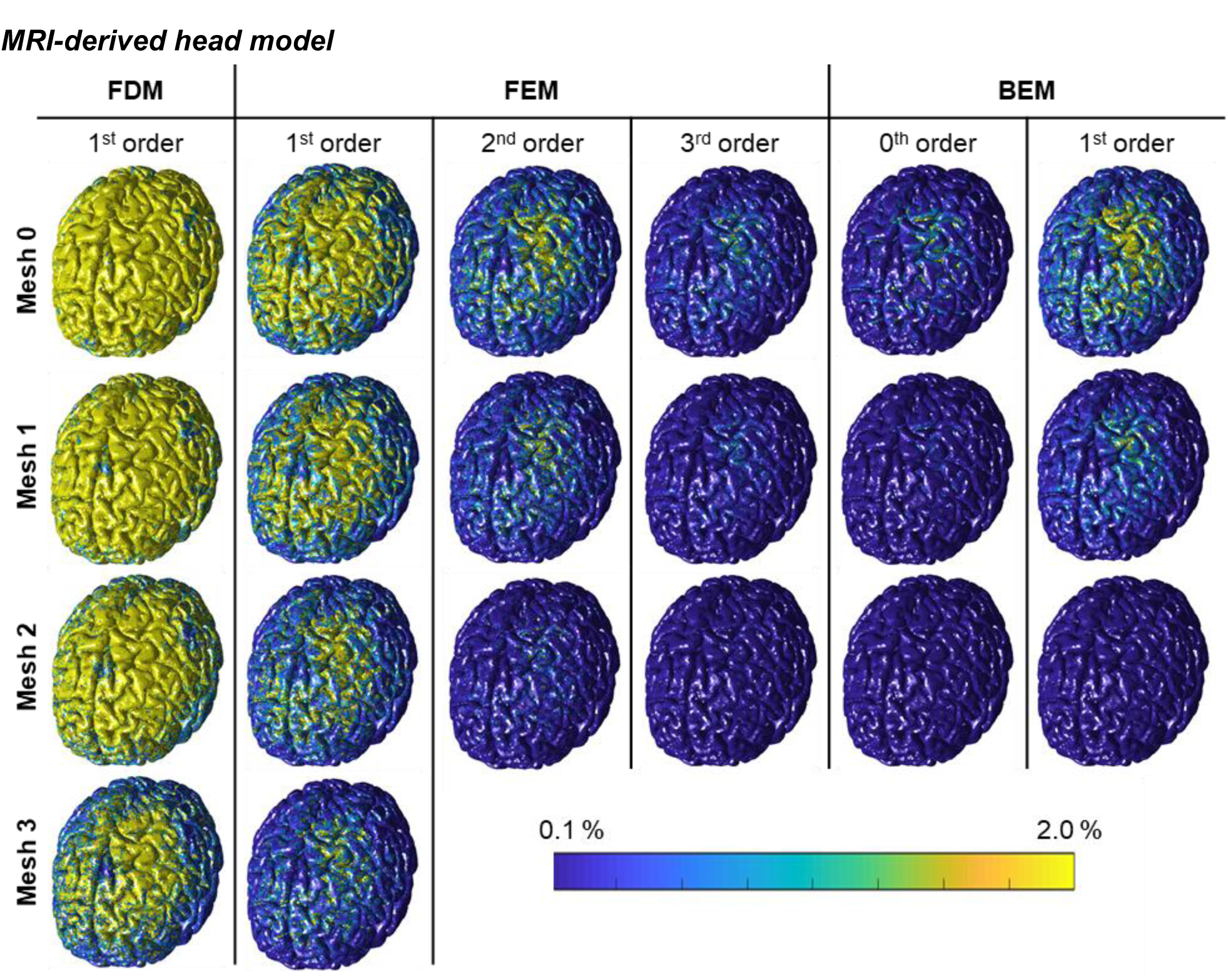
Pointwise error of cortical E-field. Results are arranged in columns by method. Rows are arranged with increasing number of mesh refinements top to bottom.

The L^2^ norm errors are shown in Fig. 4. From least to most accurate: FDM, 1^st^ order FEM, 2^nd^ order FEM, 1^th^ order BEM, 3^rd^ order FEM, and 0^st^ order BEM. The accuracy of FDM lags the accuracy of 1^st^ order FEM by one refinement. Both FDM and 1^st^ order FEM lag all other methods in accuracy by at least two refinements. 2^nd^ order FEM has an error that is nearly twice that of 1^st^ order BEM. Like the inhomogenous sphere case, 1^st^ order BEM, 0^th^ order BEM, and 3^rd^ order FEM are marginally different in error.

**Figure 4.**
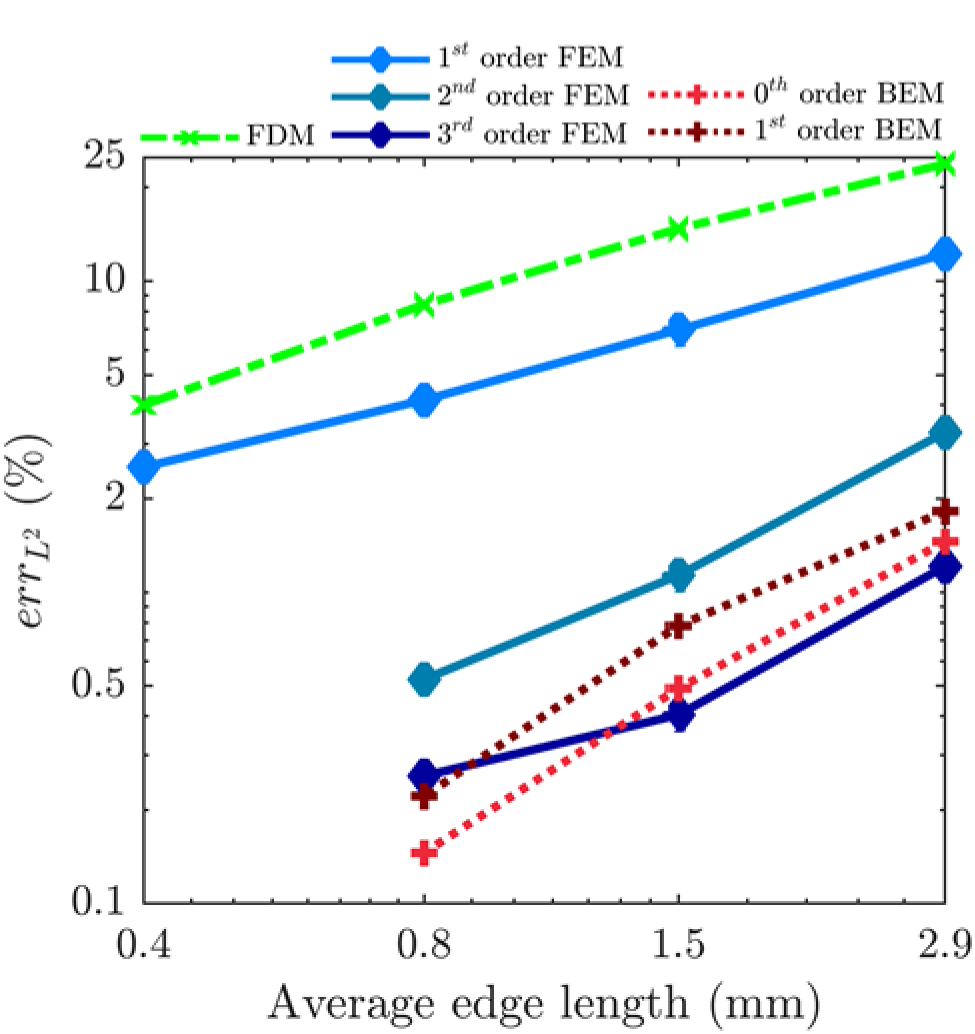
Global L^2^ norm error of cortical E-field in MRI-derived head model.

Note that the generation of the BEM system of equations requires the approximation of integral by quadrature, and the achieved error is dependent on the fidelity of this approximation. Whereas prior studies used 1^st^ order quadrature [29,43], we optimized quadrature orders for accuracy (for details see the supplement). Results given in the supplement indicate that 1^st^ order FEM has an error about 1–1.4 times higher than BEM with 1^st^ order quadrature [29,43].

Finally, to compare the computational demands, each simulation was run using a single core of an Intel(R) Xeon(R) Gold 6154 CPU computer node with 3.0 GHz 36 core processor and 768 GB of memory. Figure 5 shows the CPU runtime for each of the methods. We divided the CPU runtime into two phases: the setup phase, where a linear system of equations is assembled, and the solution phase, where the system of equations is solved iteratively. For FEM methods, increasing the order or refining the mesh result in similar CPU runtime increases. For BEM methods, the setup phase of the 1^st^ order BEM requires roughly three times the CPU runtime relative to 0^th^ order BEM, and the solution phase requires nearly the same amount of CPU runtime for both methods. The CPU runtimes of all methods could be significantly reduced by using lower level languages and parallelizing the computational routines. For example, we observed a tenfold reduction of the CPU runtime of the solution phase of BEM using shared-memory parallelism with ten cores. More detailed discussion of CPU runtime and possible speed-ups is given in the supplement. FEM and BEM implementations are significantly different and across solver comparisons would require optimizing both implementations. Independent of implementation, with each mesh refinement the CPU runtime for BEM increases by a factor of 4, whereas that of the FEM increases by a factor of 16. As a result, BEM implementations that leverage FMM are asymptotically superior to FEM in terms of CPU runtime.

**Figure 5.**
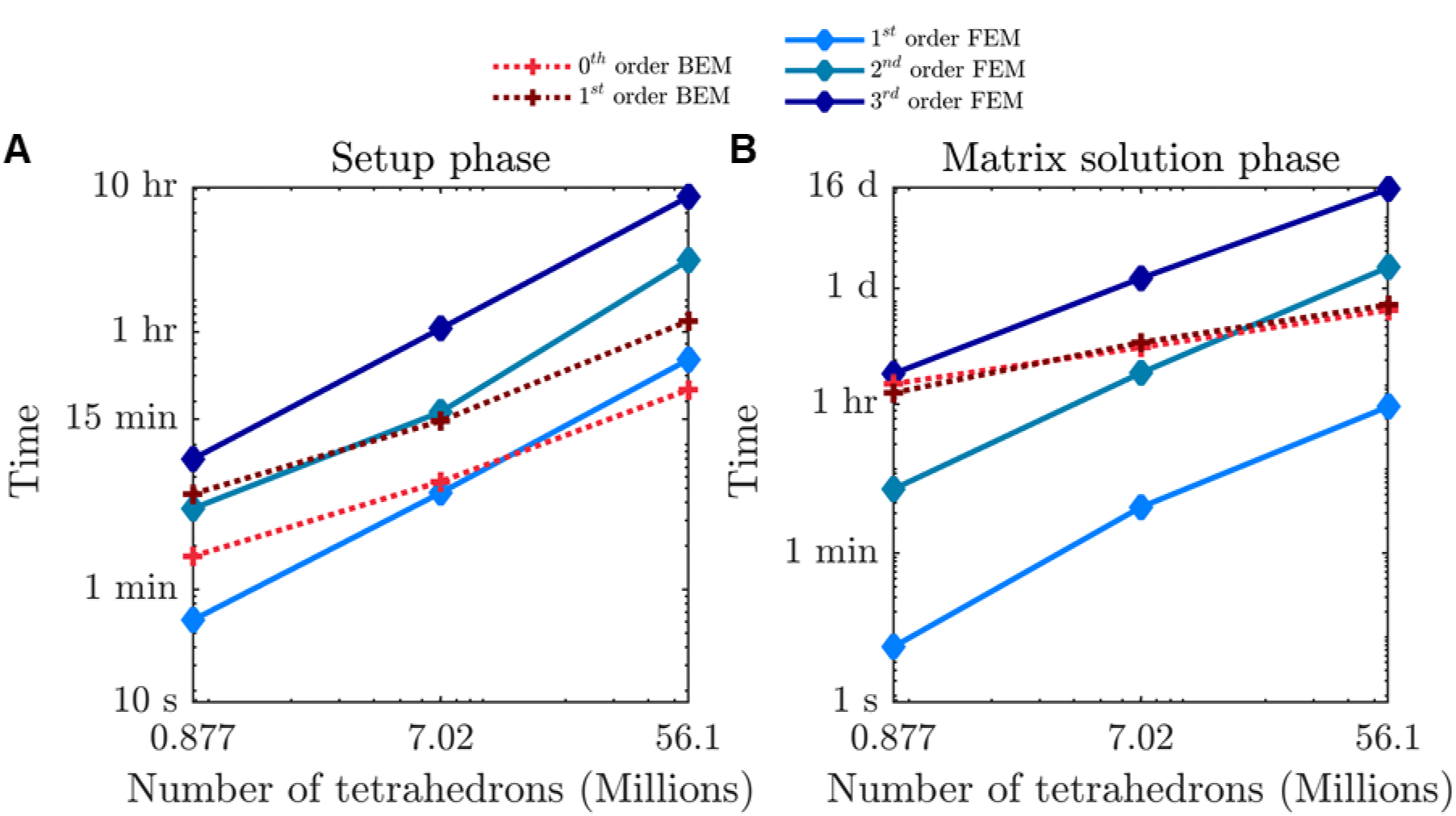
CPU runtimes of simulations with MRI-derived head model.

### Coil modeling comparison

We consider the error of the primary E-field resulting from numerical errors in the current modeling for the coil with thick wire cross-section. As reference we use a high-resolution simulation result from the coil discretized with 12,124,160 tetrahedron elements having an average edge length of 0.5 mm. The L^2^ norm and maximum pointwise error of the primary E-field induced in the spherical head model for each coil winding discretization is shown in Fig. 6. The magnetic dipole approach is more efficient than the current discretization approach, as it requires approximately an order of magnitude fewer dipoles for comparable error levels. Nevertheless, the magnetic dipole approach used here does not account for gaps between coil turns, and the L^2^ error stagnates near 0.2%. Modeling fine geometrical details and more complicated coils requires partitioning the flux integral domain, which is not always straightforward. For the current discretization approach, the primary E-field can be determined directly using a computer aided design model of the coil. Thus, the current discretization approach provides flexibility at the cost of slightly increased computation. This cost can be virtually removed by precomputing the primary E-field and storing it [11] or using FMM methods. To achieve errors below 2%, it is recommended to use > 3,000 uniformly distributed current dipoles or > 200 uniformly distributed magnetic dipoles.

**Figure 6.**
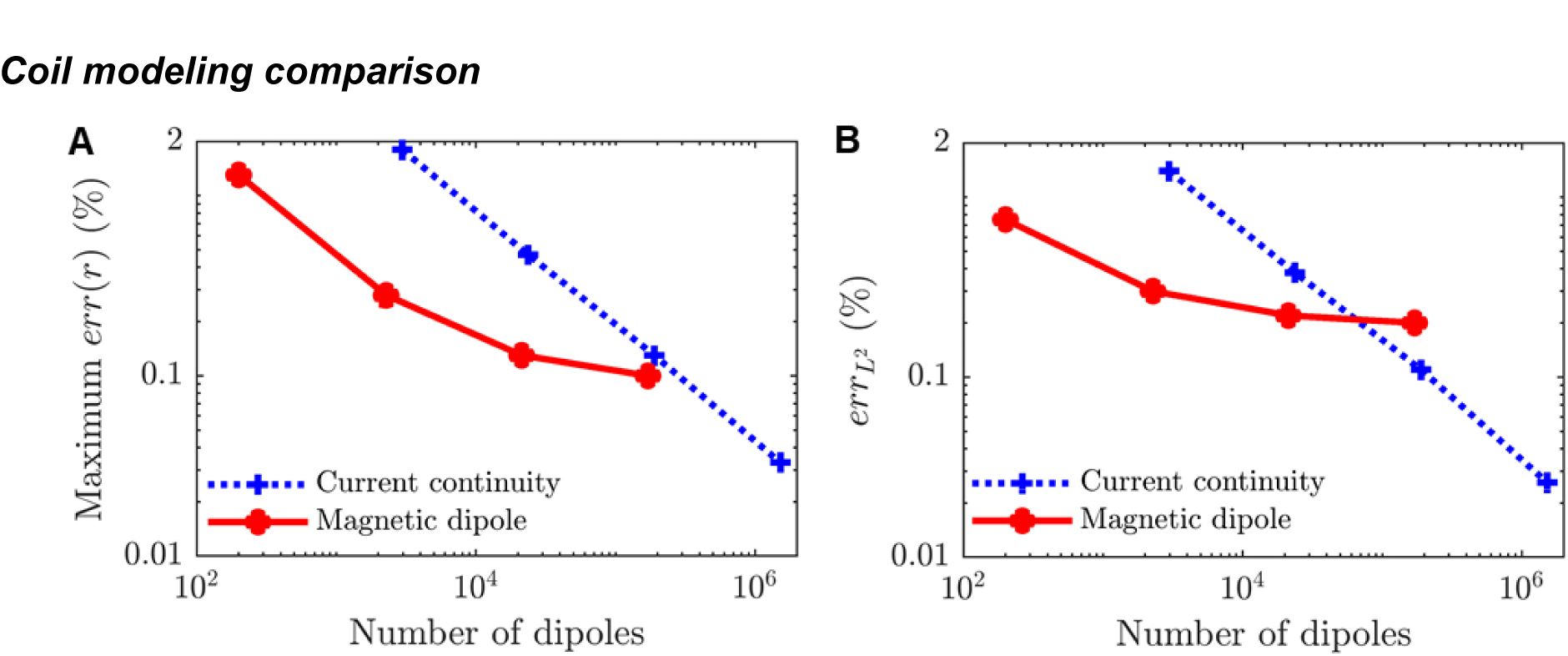
Primary E-field (A) pointwise error and (B) L^2^ norm error using current discretization and equivalent magnetic dipole approach.

Next, we consider numerical errors in the primary E-field stemming from ignoring eddy current effects in the coil current modeling. We consider the figure-8 coils with both thin (1 mm by 7 mm) and thick (1.9 mm by 6 mm) cross-section wires at a frequency of 3 kHz and 6 kHz. The currents along a coil cross-section are shown in Fig. 7. There is significant current redistribution along the vertical direction for both coils. In the horizontal direction, there is significantly more variation in the distribution of the current for the thick than thin wire coil.

**Figure 7.**
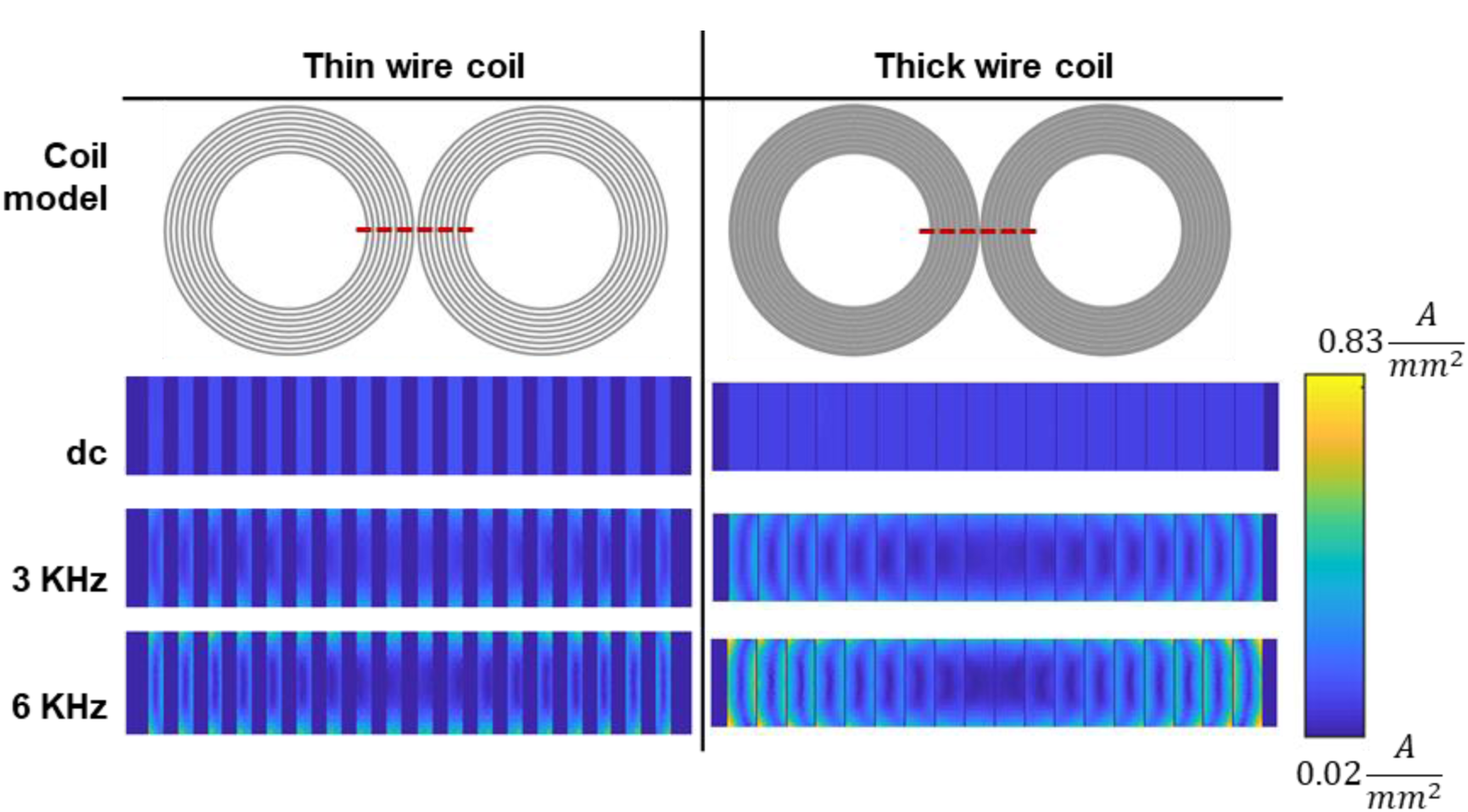
Top view of each coil model along with current distribution at dc, 3 KHz, and 6 KHz in the wire-cross section denoted by red dashed line.

The L^2^ and pointwise differences of the primary E-field sampled in the spherical model were both below 0.4% for 3 kHz and 1.0% for 6 kHz for the thin wire coil, but increased for the thick model respectively to 1.7% and 2.6% for 3 kHz, and 3.2% and 5.0% for 6 kHz. Comparing the total E-field generated in the MRI-derived head model cortex by the thick wire coil with and without eddy current redistribution revealed L^2^ and maximum pointwise differences of 1.6% and 1.9% for 3 kHz, and 3.1% and 3.8% for 6 kHz, respectively. Therefore, when the TMS pulse has frequencies beyond about 3 kHz, the introduction of eddy currents is likely important to achieve a numerical error below 2%. Finally, the primary E-fields induced by the thin and thick wire coils (without including eddy effects) differed by more than 3%, confirming that accurate representation of the wire geometry is important [44].

## DISCUSSION

In TMS modeling there are three main sources of numerical error: coil modeling fidelity, mesh resolution, and accuracy of the numerical method. Most methods typically employed for modeling coil currents sufficiently suppress numerical errors. TMS coil wire usually has a small cross-section or of a litz wire, and therefore eddy current effects are typically negligible for TMS simulation. However, to attain TMS coil E-field errors below 2%, modeling eddy currents might be necessary in extreme cases when the wire is made of a solid conductor with a relatively large cross-section, and/or very brief pulses are used.

The mesh resolution mediates both how well the mesh approximates the tissue geometry and how accurate the FEM/BEM solution is. FDM meshes typically do not conform well to tissue boundaries, resulting in additional errors relative to FEM and BEM methods. Head models with spherical geometry cannot be used to estimate the numerical errors in a realistic TMS setup; for a given mesh resolution, the numerical errors even in the inhomogeneous spherical geometry are at least a factor of ten smaller than in a realistic head geometry. Unsurprisingly, 1^st^ order FEM methods converge slowly with respect to increasing mesh resolution. This is because they employ a piecewise constant approximation of the E-field within each mesh element. Previous attempts of using adjoint double layer BEM formulation to analyze TMS-induced E-fields [29,43] exhibit errors similar to those of 1^st^ order FEM methods (see supplement). Increasing the order or refining the mesh of FEM results in similar CPU cost increases. Increasing the order in FEM reduces the error more rapidly than increasing the mesh resolution. 2^nd^ and 3^rd^ order FEM, and 0^th^ and 1^st^ order BEM exhibit similar levels of accuracy near tissue interfaces. This lack of improvement through P-refinement of the BEM methods could be due to excessive regularity imposed through the use of piece-wise linear basis functions, as conforming discretizations of the adjoint BEM operator admit piece-wise discontinuous functions [26]. Because of the complicated cortical geometry, a dense mesh is required to sufficiently resolve all its geometric features. As such, the use of a 2^nd^ order FEM or 0^th^ order BEM method may diminish the numerical error to acceptable levels without the need of additional mesh refinements in practical applications.

## CONCLUSIONS

We benchmarked various methods for simulating TMS induced E-fields. Whereas at present 1^st^ order FEM is most commonly used, 2^nd^ order FEM or 0^th^ order BEM appear best for achieving negligible numerical error relative to modeling error, while maintaining tractable levels of computation. FDM requires excessive mesh densities to result in accurate cortical boundary representation, limiting the computationally achievable accuracy. In addition to the numerical errors arising from the solver, if the TMS coil windings use a large cross-section solid wire, the eddy currents within the coil windings may have to be modeled to reach error levels below 2%.

## ACKNOWLEDGEMENT

Research reported in this publication was supported by the National Institute of Mental Health and the National Institute of Neurological Disorders and Stroke of the National Institutes of Health under Award Numbers RF1MH114268 and R01NS088674-S1. The content of current research is solely the responsibility of the authors and does not necessarily represent the official views of the National Institutes of Health.

## SUPPLEMENTARY MATERIAL

### FEM solver

The TMS coil is driven by coil currents **J**(**r**^′^;*t*) = P(t)**J**(**r**^′^), where P(t) is the pulse-waveform, and **J**(**r**^′^) is the spatial distribution of the current. At the low frequencies used for TMS, quasi-stationary assumptions are valid, and the temporal variation and spatial variation of the E-field are separable (i.e.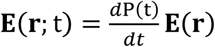., where **E**(**r**) is solely dependent on **J**(**r**^′^) [45]. The spatial variation of the E-field induced in the head **E**(**r**) is the sum of the incident (primary) field **E**_p_(**r**) due to the coil currents and a secondary contribution **E**_s_(**r**) = −∇ϕ(**r**) from the scalar potential ϕ(**r**), where

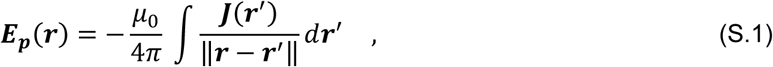

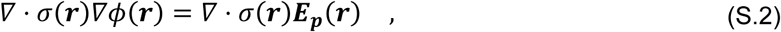

μ0 is the electric permeability of free-space, and σ(**r**) is the tissue conductivity. Furthermore, the normal component of the E-field on the surface of the head is zero. To solve for ϕ and ∇ϕ, first, the head model is approximated by a mesh consisting of tetrahedrons, and each tetrahedron is assigned a constant tissue conductivity. The scalar potential ϕ is approximated by

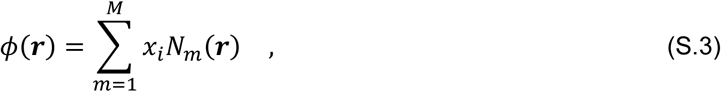

where *N_m_*(**r**) are piecewise linear, quadratic, and cubic nodal elements for 1^st^, 2^nd^, and 3^rd^ order FEM, respectively [17], and **x** = (*x*_1_, …, *x_M_*) are expansion coefficients. A linear system of equations in terms of **x** as unknown is assembled by a standard Galerkin procedure, i.e. each equation is generated by multiplying Eq. S2 by a testing function *N_m_*(**r**), and integrating it over its support. This results in

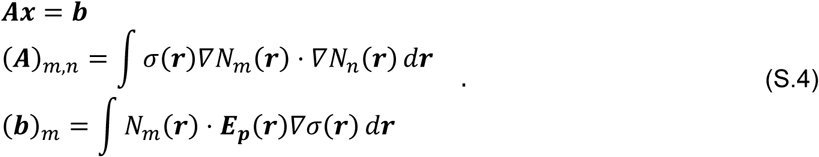

To evaluate (**A**)*_m,n_* exact values of integrals in terms of tetrahedron vertex locations provided in [17] are used. To determine (**b**)*_m_*, a 2^th^, 3^rd^, and 4^th^ order accurate quadrature rule is used for 1^st^, 2^nd^, and 3^rd^ order FEM, respectively. All codes use quadrature rules [40]. The linear system of Eqs. (S.4) is solved using a minimal residual (MINRES) iterative solver to a relative residual of 10^−7^ [46].

### FDM solver

The FDM solver also solves Eq. S.2. The main difference is that the head model is approximated by a regular mesh consisting of identical cuboid elements. Furthermore, first order nodal elements are used to approximate the scalar potential [18].

### BEM solver

An implementation of the BEM method [29] assumes that the scalar potential arises from charges ρ(**r**) on tissue interfaces. This charge generates a scalar potential

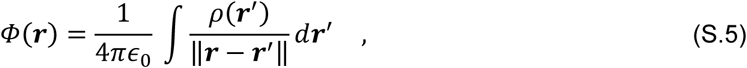

and secondary E-field

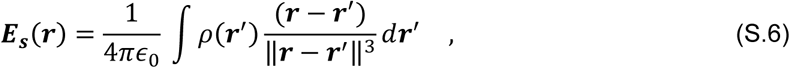

where ϵ0 is the permittivity of free space. Current continuity across a tissue interface dictates that

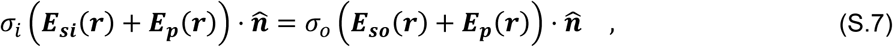

where *σ_i_* is the inner and *σ_o_* is the outer tissue conductivity, **E**_si_(**r**) and **E**_so_(**r**) is the secondary E-field an infinitesimal distance interior and exterior to the interface, respectively, and 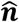 is the tissue interface normal pointing towards the outer tissue. If the tissue interface is smooth, this results in the following integral equation

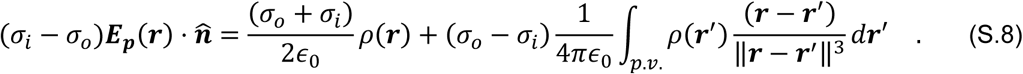

Here the residue of the integral of Eq. S.4 has been extracted as 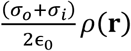., and p.v. is used to denote the principal value.

To solve Eq. S.8 for the charge, tissue boundaries are approximated by meshes consisting of triangles. Then, the charge is expanded as

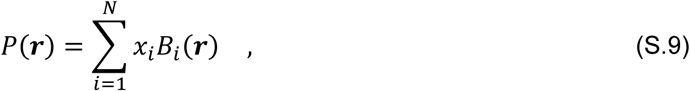

where *B_i_*(**r**) are pulse functions for 0^th^ order BEM and *B_i_*(**r**) are triangular piecewise linear nodal elements for 1^th^ order BEM [41]. A linear system of equations in terms of **x** = (*x*_1_, …, *x N*) as unknown is derived by applying a standard Galerkin procedure on Eq. S.8. The resulting system of equations is

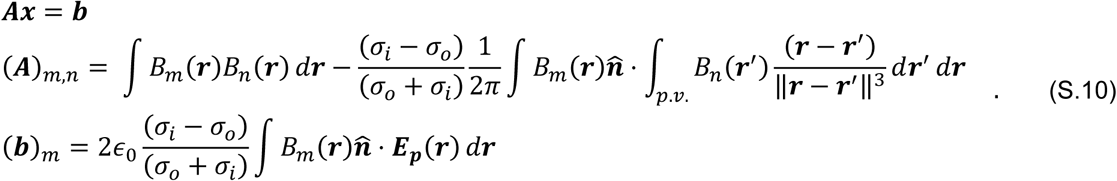

The first term of (**A**)*_m,n_* is computed exactly. For the second term, the quadrature rule used depends on the distance between the inner integration and outer integration triangular support center. Specifically, we distinguish between entries corresponding to near triangles and far apart ones. Two triangles are considered to be near to each other if they are a distance less than *d_near_* apart, measured from triangle centers, or they share a vertex. Otherwise, they are considered far apart. For near triangle entries, we compute the outer integral using an 8^th^ order accurate rule and the inner integral exactly. For far apart triangle entries, a 2^nd^ order accurate quadrature rule is used [40]. To lower computational costs, the FMM library with precision set to δ is used to compute all far apart triangle entries on-the-fly each time the matrix is applied to a vector. The linear system of Eqs. S.10 is solved using TFQMR [46] iterative solver to a relative residual of relres.

We selected the values for *d_near_*, quadrature orders, and δ as follows. (i) We ran all sphere numerical experiments using *d_near_* equal to 5 times the maximum edge length, 8^th^ order quadrature integrals, and δ = 5 × 10^−10^. (ii) For near triangle terms, we decreased the quadrature order to as low as possible without increasing error. (iii) For near triangle terms, starting from 12^th^ order accurate outer integral of nearby terms, we lowered the quadrature rule, and determined that the accuracy degrades below 8^th^ order. This is not surprising since there is a logarithm in the analytical expression for the inner integral [47,48] that goes to infinity as one approaches an observation point on the edge of the triangular source distribution. Although this logarithmic singularity is integrable, it still increases the error for nearby elements and higher order integration rules are essential [32]. (iv) We decreased the value of *d_near_* to as low as possible without increasing error levels. (v) We re-ran the simulations each time decreasing δ and chose the minimum possible without increasing error. For the MRI-derived head model, we used the same parameters as for the inhomogenous sphere model. To ensure these parameters translated to the MRI-derived head model, we ran all simulations with parameters δ = 0.5 × 10^−9^, 4^th^ order quadrature rule, and *d_near_* = 1 × *maximum edge length*. No significant error difference was observed. Values used for parameters *d_near_*, quadrature orders, δ, and relres are summarized in Table S.1. The homogenous sphere simulations have a lower level for δ because otherwise the L^2^ error levels of the E-field would have stagnated with increasing mesh resolution at 10^−4^; however, above that they were the same as with the lower threshold of 5 × 10^−3^. The same was observed with parameter *d_near_*, which we had to increase to 1.8 to reach L^2^ error levels below 10^−4^. If we had used the same parameters as with the inhomogeneous sphere, the maximum pointwise E-field errors for all results would have remained unchanged. This is important because the maximum pointwise E-field error is more indicative of local error on regions of interest than L^2^ error. As such, using the set of simulation parameters optimized for the inhomogenous sphere model is likely warranted to maintain computational tractability for general simulation frameworks.

For consistency we chose relres = 10^−7^ for all FEM and BEM simulations. The accuracy of the resulting **x** is such that ‖**x**‖ ≤ C × relres [49], where C is the condition number of **A**. Double layer potential formulations are known to have a low-condition number relative to FEM for the same discretization because of their identity plus compact operator form [50]. The relres parameter can likely be increased beyond what was used here while maintaining the same accuracy level on the E-field.

To determine the E-field from a charge distribution, if a source distribution was less than *d_near_* from the observation point, then its contribution was computed analytically, otherwise we used FMM (δ = 5 × 10^−10^) along with a 6^th^-order accurate quadrature rule.

**Table S.1.** Table S.1. BEM solver parameters used for simulations.

| Head model | Homogenous sphere |  | Inhomogeneous sphere |  | MRI-derived |  |
| --- | --- | --- | --- | --- | --- | --- |
| BEM order | 0<sup>th</sup> | 1<sup>st</sup> | 0<sup>th</sup> | 1<sup>st</sup> | 0<sup>th</sup> | 1<sup>st</sup> |
| $d_{\text{near}}$<br>maximum edge length | 0.5 | 1.8 | 0.5 | 0.5 | 0.5 | 0.5 |
| Quadrature order | 2<sup>nd</sup> | 2<sup>nd</sup> | 2<sup>nd</sup> | 2<sup>nd</sup> | 2<sup>nd</sup> | 2<sup>nd</sup> |
| $\delta$ | $5 \times 10^{-3}$ | $5 \times 10^{-7}$ | $5 \times 10^{-3}$ | $5 \times 10^{-3}$ | $5 \times 10^{-3}$ | $5 \times 10^{-3}$ |
| relres | $10^{-7}$ | $10^{-7}$ | $10^{-7}$ | $10^{-7}$ | $10^{-7}$ | $10^{-7}$ |

### Coil eddy current solver

Assume the coil is driven with a time-harmonic current source with amplitude *I_in_* flowing into port A and out of port B. (In what follows, the temporal variation exp(*iωt*), where *i* is the imaginary unit, is assumed and suppressed for notational brevity.) The current continuity dictates that

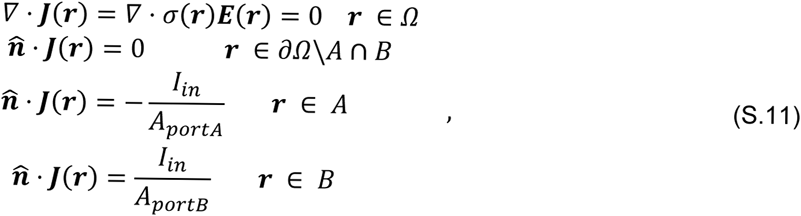

where Ω is the coil wire, ∂Ω\A ∩ B is the coil wire boundary excluding A ∩ B, A is port A with area *A_portA_*, B is port B with area *A_portB_*, and 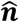 is an outward pointing normal. Furthermore, the total E-field inside the coil is due to the coil conduction currents **J**(**r**) = σ(**r**) **E**(**r**), i.e.

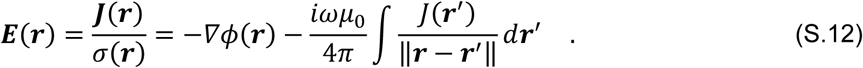

Here we assumed that the current source forms a complete loop, thereby, not generating a scalar potential contribution. We expand the current in terms of a dc dissipative part −σ(**r**)∇∇(**r**) and rotational part ∇ × **W**(**r**) as **J**(*r*) = −σ(**r**)∇*ϕ*(**r**) + ∇ × **W**(**r**). Equations S.11–S.12 become

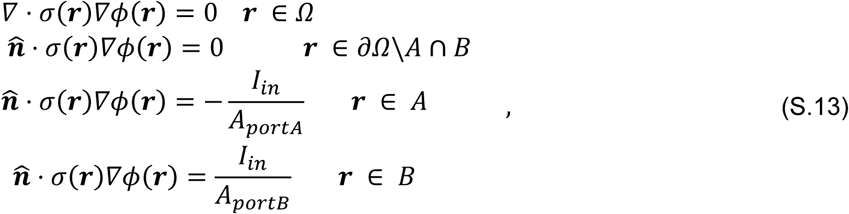

and

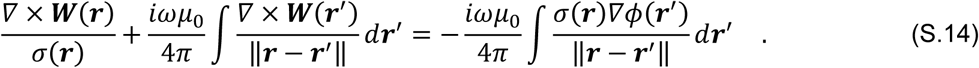

We solve Eqs. S.13–S.14 by first approximating the coil winding as a mesh consisting of tetrahedral elements. Then, we solve Eq. S.13 using the method described in the FEM solver section. The resulting current flows uniformly through the conductor and is the current that would flow assuming no temporal variation, i.e. dc current. To incorporate eddy current effects, we expand ∇ × **W**(**r**) using a loop basis [51], which is derived from the curl of 1^st^ order edge elements as defined in [17]

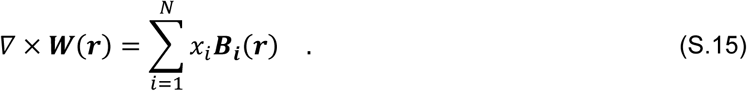

Here **B**_i_(**r**) is a loop basis function corresponding to the ith edge and **x** = (*x*_1_, …, *x N*) are the expansion coefficients. Because the normal component of the current at the boundary is zero, we do not include boundary loop edges in the expansion. Applying a standard Galerkin procedure to Eq. S.14 results in a linear system of equations

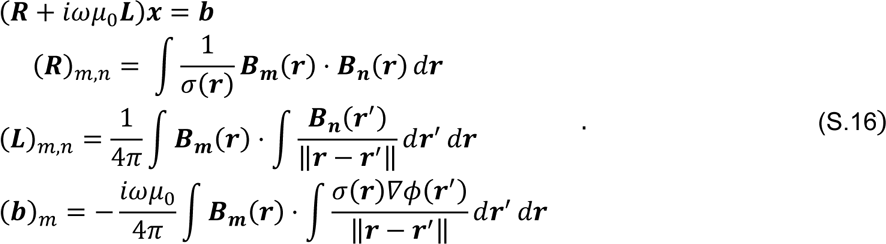

We compute (**R**)*_m,n_* using relations available in [17]. The inner integral of (**R**)*_m,n_* is computed exactly for elements less than a distance *d_near_* using formulas in [52], and using a 2^nd^ order quadrature rule for all other elements. For all our simulations, *d_near_* is measured from tetrahedron centers and is set as the maximum edge length of the tetrahedron mesh. The outer integral is also computed using a 2^nd^ order quadrature rule. To save computation time, the FMM library with precision set to 10^−8^ is used to compute multiplications of a vector with **L**. To determine (**b**)*_m_*, again a 2^nd^ order quadrature rule is used.

The matrix **R** is a Gramm-matrix of a loop basis set and is well known to have an unbounded ratio of maximum to minimum nonzero singular values [53], which results in slow convergence when solved using iterative solvers. However, applying **R** requires orders of magnitude less computational resources than to apply **R**. To speed-up the computation we solve a modified system of equations

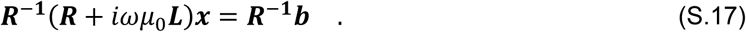

Here to compute **R**^−1^_y_ we solve the following system **Rz** = **y** for **z** iteratively using MINRES [46] to a relative residual of 10^−6^. System of Eqs. S.17 is iteratively solved using GMRES to a relative residual of 10^−6^, and it typically takes 10–20 iterations to converge.

### Comparison of Galerkin BEM with delta-testing BEM

We compared the delta-testing BEM discretization used in [29,43] with the Galerkin-testing BEM used in this work. In summary, the delta-testing scheme requires less computational resources but is less accurate than the Galerkin BEM. As such, the optimal choice between the Galerkin and delta-testing schemes depends on the accuracy level required by the user.

The delta testing scheme [29,43] used 0^th^ order basis elements and delta-testing functions. Briefly, for near triangle entries, the inner integral is computed analytically, and the outer integral and all other integrals are computed using single point quadrature (i.e. delta testing). In [29], the value of *d_near_* used depends on the simulation results, and it is in the range of 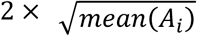 to 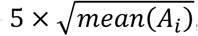, where *mean*(*A_i_*) is the mean area of the mesh triangles. In [43], *d_near_* is set to zero and only entries that correspond to neighboring triangles, i.e. triangles sharing a node, are considered as near elements. Based on this we have developed three formulations: *d_near_* set to 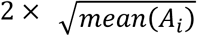, *d_near_* set to 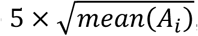, or only including neighboring triangles. Finally, the system of Eqs (S.10) is solved to *relres* = 10^−4^, and the total E-field is computed from the charge using a single point quadrature. In [43], a least squares error metric 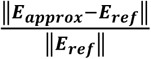 is used, where *E_approx_* and *E_ref_* are vectors of E-field samples on a spherical shell and ‖⋅‖ denotes Euclidian distance. In what follows we compute the above metric for spherical head models to show equivalence between the delta-testing BEM implementation and the one used in [43]. (The least squares error is the discrete equivalent of the L^2^ norm error; for dense enough sampling the two converge. In the main manuscript, we did not include least squares error as a metric because it would be redundant with L^2^.)

First, we compare the delta-testing BEM implementation with 1^st^ order FEM and Galerkin BEM implementations on the inhomogenous spherical head model. In Fig. S.1A we show the least squares error of the E-field on a shell 5 mm into the head like in [43]. For the delta-testing BEM discretizations, we achieved least squares errors of 2–2.9% and 0.9–1.2% with an average edge length of 4.8 and 2.8 mm, respectively. In [43], errors of 2.7% and 1.6% were observed for an average edge length of 4.2 and 2.9 mm, respectively. Compared with the SimNIBS software[11], the approach in [43] uses a different interpolation scheme termed super-convergent patch recovery[54], resulting in errors of 6.5% and 6.1% for a mesh having average edge length of 4.2 and 2.9 mm, respectively. We observed a 3.7% and 2% error for an average edge length of 4.8 and 2.8 mm, respectively. There are small differences in the errors we observe, as expected for different sphere meshes and sampling points.

What follows only pertains to meshes having average edge length below 7 mm, and the best of the three implementations of the delta-testing BEM. We exclude other results because the triangles are large and the one-point quadrature rule used in the delta-testing scheme to compute the E-field from the charges results in erroneous E-field values; furthermore, the large edge lengths are not representative of what is used in practice. The maximum error on the spherical 77.5 mm radius shell is shown in Fig. S.1B. The maximum error of the 1^st^ order FEM is 2.3–8.4 times higher than that of the delta-testing BEM. The delta-testing BEM has a maximum error 4–10 times higher than the Galerkin BEM. The L^2^ norm error is shown in Fig. S.1C. The 1^st^ order FEM has 1.5–1.8 times the error of the delta-testing BEM. The delta-testing BEM implementation has an error of 74–127 times that of the Galerkin BEM.

**Figure S.1.**
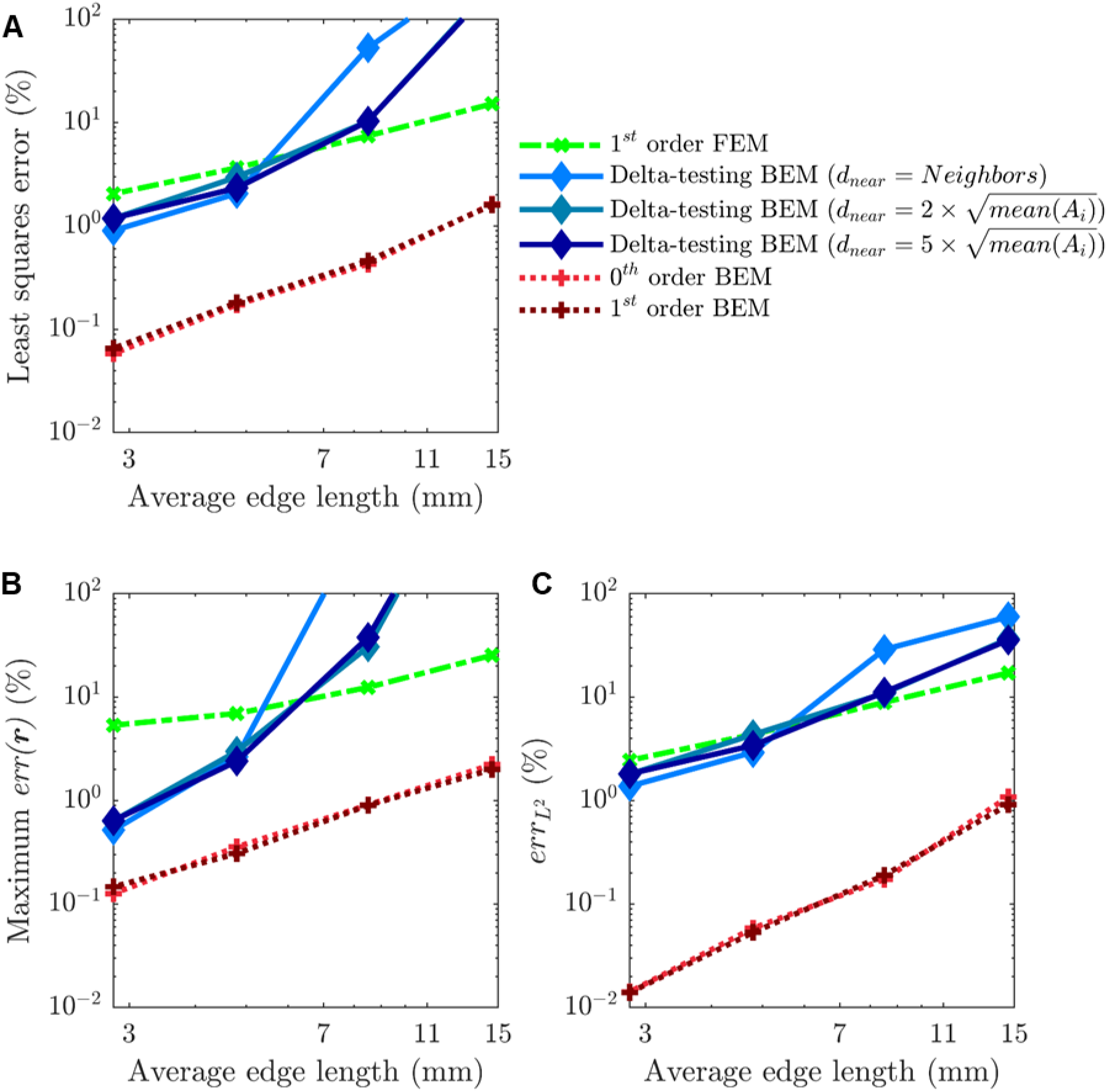
Error results for the inhomogenous sphere head for delta-testing BEM and Galerkin BEM. 1^st^ order FEM and Galerkin BEM results in B and C are reproduced from Figures 2B and 2D, respectively.

The pointwise errors 0.5 mm into the head are shown in Fig. S.2. For Mesh 0, the pointwise errors are better for 1^st^ order FEM than the delta-testing BEM. For other meshes, the delta-testing BEM outperforms 1^st^ order FEM. Again, pointwise error is lower with Galerkin BEM than other methods. The L^2^ norm errors are shown in Fig. S3. For the original MRI-derived mesh, the 1^st^ order FEM and delta-testing BEM have approximately the same accuracy. For other meshes, the 1^st^ order FEM has an error that is 30%–40% higher than that of the delta-testing BEM. Again, Galerkin BEM methods outperform the others in terms of accuracy.

**Figure S.2.**
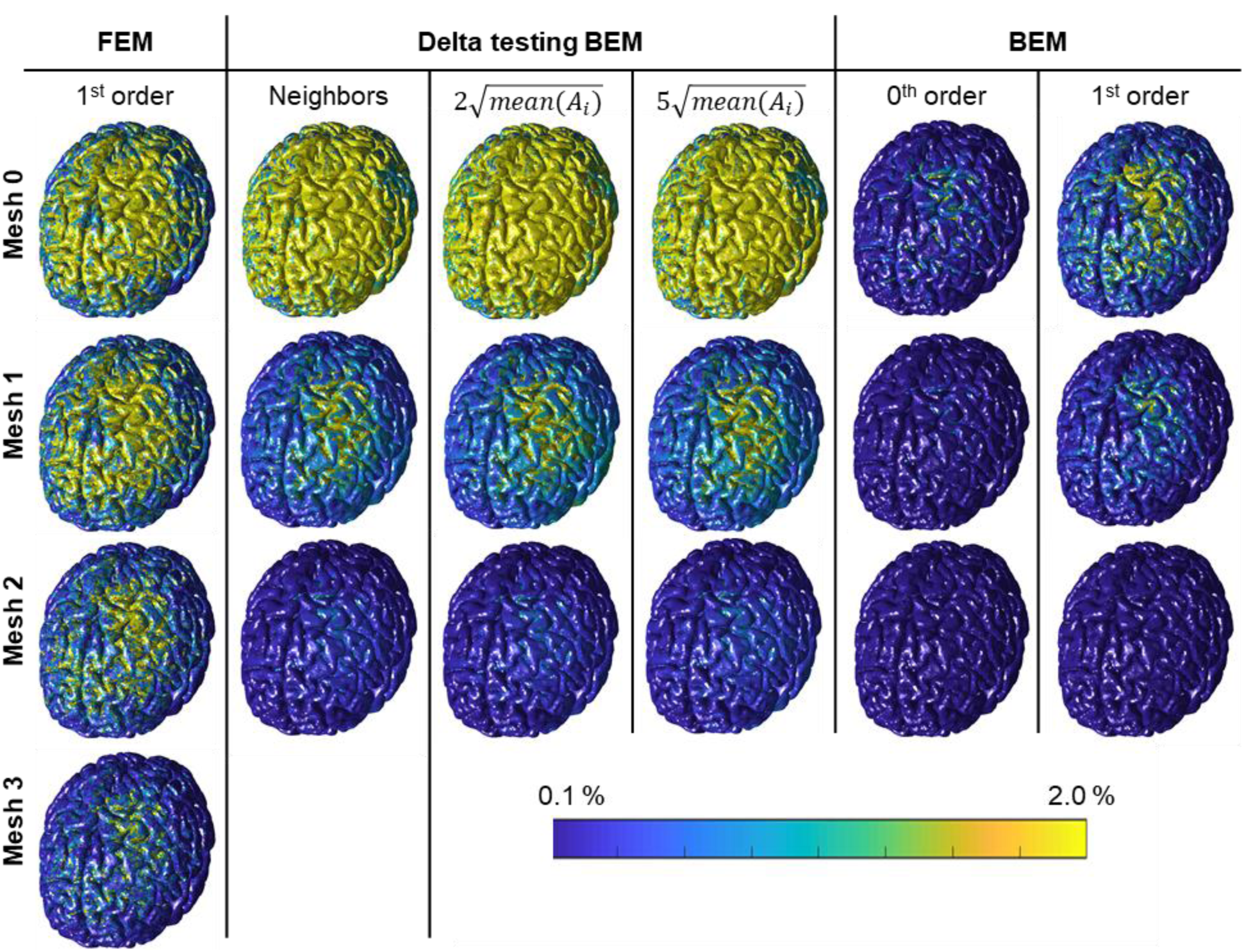
Pointwise error of cortical E-field in MRI-derived head model. Results are arranged in columns by method. Rows are arranged with increasing number of mesh refinements top to bottom. 1^st^ order FEM and Galerkin BEM results are reproduced from Fig. 3.

**Figure S.3.**
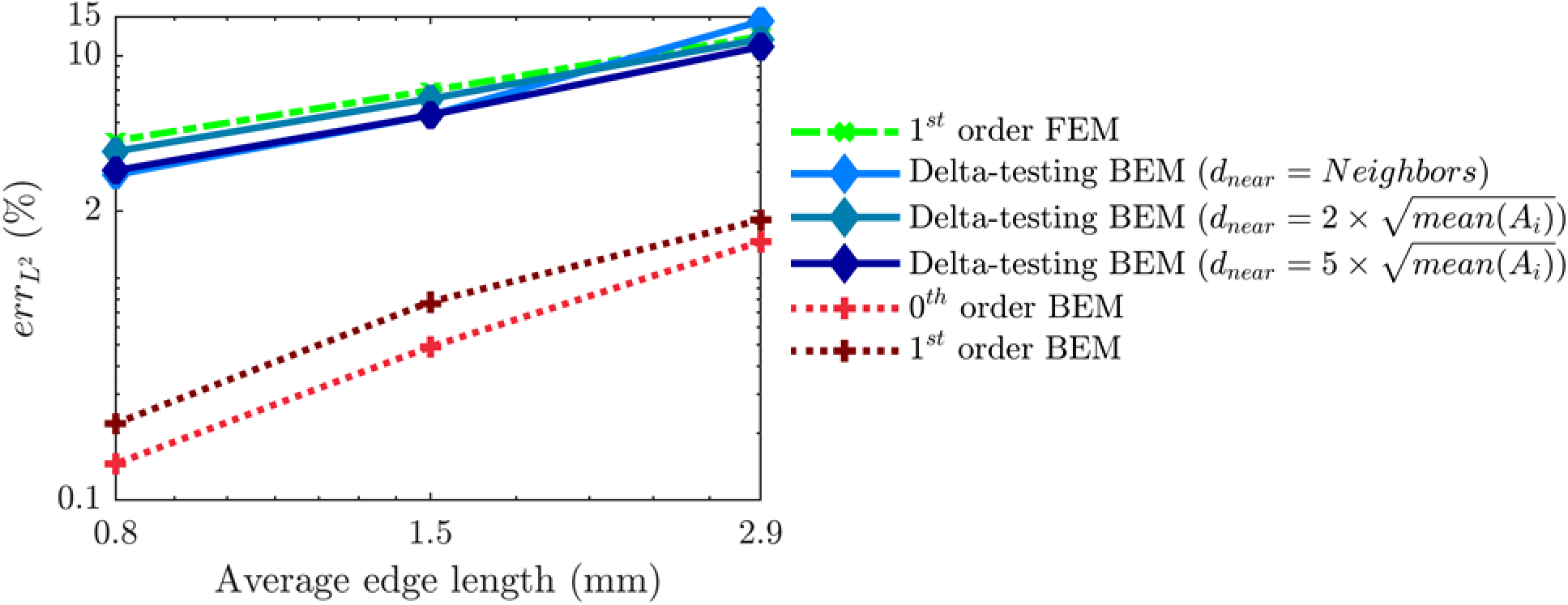
Global L^2^ norm error of cortical E-field in MRI-derived head model. 1^st^ order FEM and Galerkin BEM results are reproduced from Fig. 4.

Thus, the Galerkin BEM is more accurate than the delta-testing BEM. However, this additional accuracy is gained at the cost of additional computational resources. The near elements were computed using an 8^th^ order quadrature rule, which requires 16 points per triangle. This pertains only to the setup phase of the code, which only has to be done once and does not need to be repeated each time the solver is run for a different coil position. For the MRI-derived head model, we observed that Galerkin BEM required 2.4–6.4 times the setup time of the delta-testing BEM. When we increased the number of quadrature points per triangle from 1 to 3, Galerkin BEM required 1.8-4.3 times the solution time of delta-testing BEM. A more detailed CPU-runtime study is given in the next section.

### CPU resources for BEM and FEM

In this subsection, we present extended CPU runtime results. We only present results for the MRI-derived head, as they are more representative of typical uses of the solvers. Furthermore, since we solve each system of equations via an iterative method, we split the CPU runtime into setup phase and a matrix vector multiplication phase. For a fair comparison, all results were run on a single processor of an Intel(R) Xeon(R) Gold 6154 CPU computer node with 3.0 GHz 36 core processor and 768 GB of memory.

Fig. S.4 shows the CPU runtime for the FEM methods. The CPU runtime for the setup phase (Fig. S.4A) can be lowered significantly by porting the code to a lower level language, and using shared memory parallelism. The solution time is the cost per matrix vector multiplication (Fig. S.4B) times the number of iterations (Fig. S.5). The solution time typically dominates the setup time in FEM because the number of iterations required for convergence are typically large, ranging from 500–14,000. This cost could be lowered by choosing a higher level for relative residual error convergence criteria, which might result in a less accurate solution, or using preconditioning techniques to speed up the convergence. For easy estimation of CPU runtime for each given convergence criteria, the relres values are shown in Fig. S.5. Alternatively, the matrix vector multiplication time can be sped up. Sparse matrix vector multiplication is a built-in Matlab function, which has been optimized for performance. Nevertheless, it can still be memory-shared-parallelized to increase computational speed. Note that classical sparse matrix data structures are not amenable to efficient parallelization, and specialized sparse matrix data structures like the ones in the Portable, Extensible Toolkit for Scientific Computation (PETSc) of Argonne labs [55] have to be used.

**Figure S.4.**
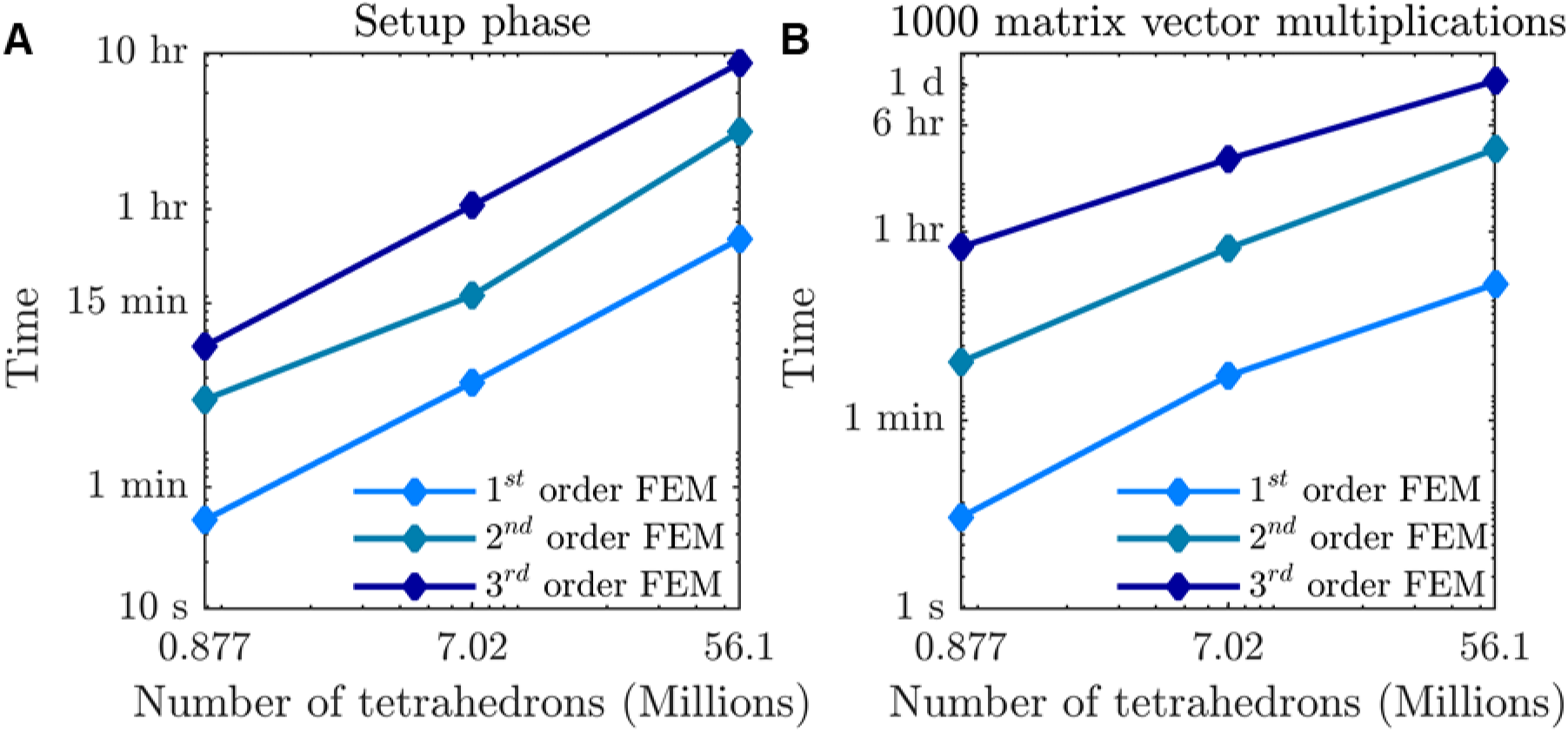
Computation times for various FEM methods versus mesh density.

**Figure S.5.**
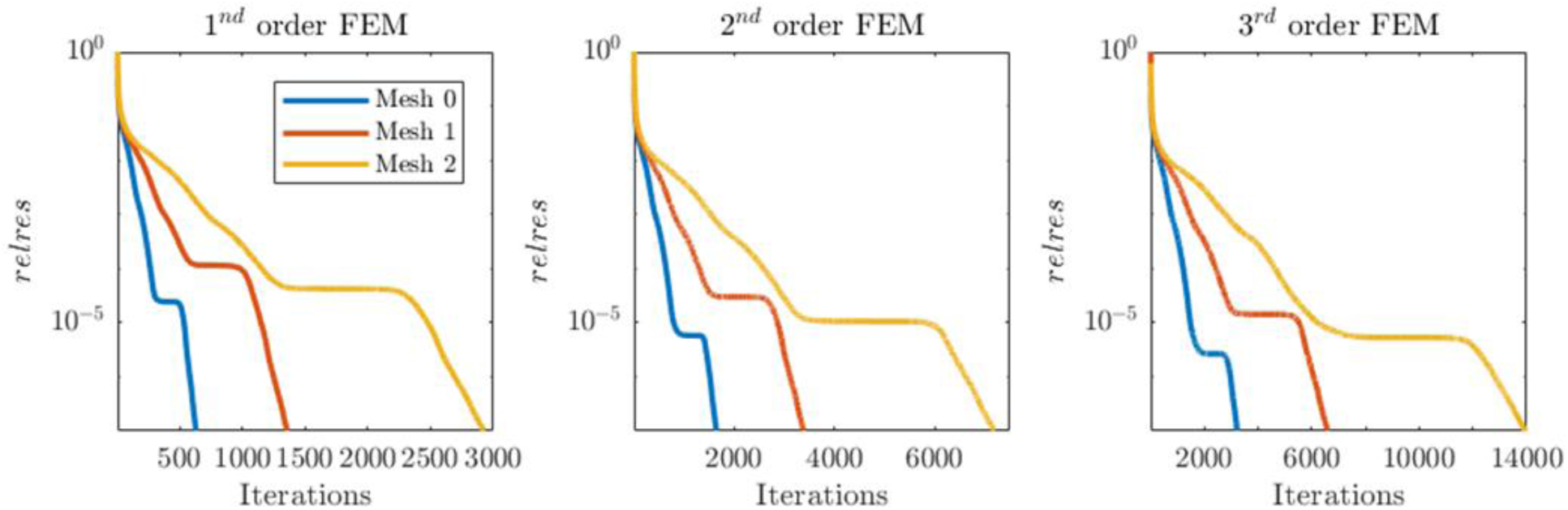
Relative residual values versus number of iterations for MINRES solution of FEM systems of equations.

Fig. S.6 shows the CPU runtime for BEM methods, with the setup phase separated into a mesh pre-processing phase (Fig. S.6A) and a near-field setup phase (Fig. S.6B). In the mesh pre-processing phase, a triangle mesh is extracted from the tetrahedral mesh and triangle mesh data structures are defined. This phase is separated from the others as most of the computation associated with it is not necessary because we can directly operate on triangle meshes without ever generating a tetrahedron mesh. In the near-field setup phase, near matrix entries are computed. Again, the CPU runtime for the near-field setup phase (Fig. S.6B) can be lowered using shared memory parallelism. For TFQMR, the solution time is two times the cost per matrix vector multiplication (Fig. S.6C) times the number of iterations (Fig. S.7). For BEM methods, the number of iterations remains relatively constant across methods and formulations. Additional results indicate that the use of GMRES lowers the required number of matrix vector multiplications by 20–30%; however, this comes at an additional cost in memory because GMRES requires the storage of iterative solution history. FMM is parallel and the cost presented here can be lowered by using memory-shared parallelism. We observed that with parallelization the CPU runtime of FMM decreases linearly with increasing number of processor cores for up 10 cores.

For FEM, each time the mesh is refined the number of tetrahedrons increases by a factor of 8, and the number of iterations double. As a result, the CPU runtime increases by a factor of 16 with each mesh refinement. This factor can be reduced to 8 using multigrid methods. For BEM, each time the mesh is refined the number of variables increases by a factor of 4, and the number of iterations remain relatively constant. The CPU runtime increases by a factor of 4 with each mesh refinement. For dense enough meshes, and independent of optimality of implementation, the BEM method presented here will always outperform the FEM method in terms of CPU resources.

**Figure S.6.**
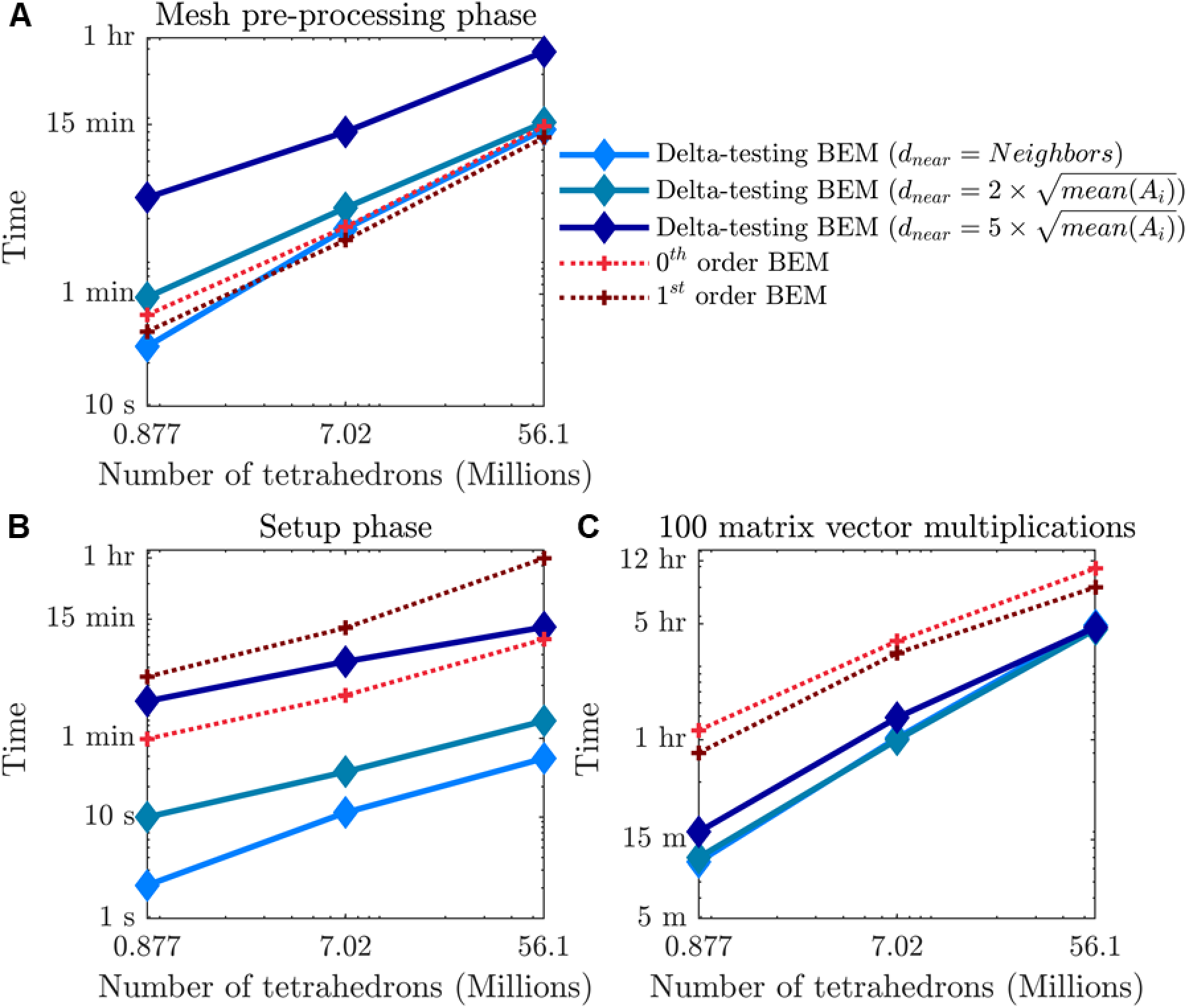
Computation times for various BEM methods versus mesh density.

**Figure S.7.**
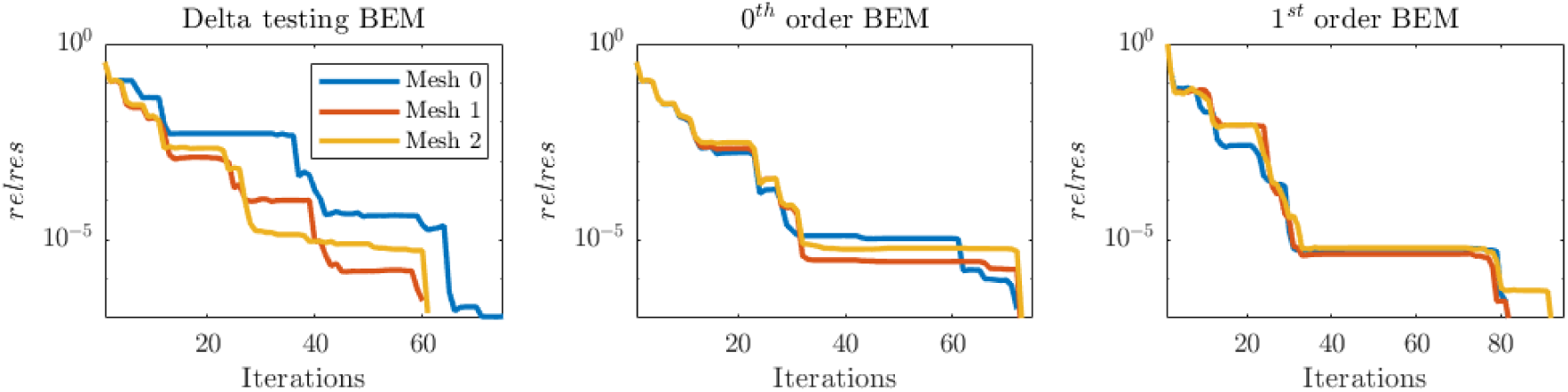
Relative residual value versus number of iterations for TFQMR solution of BEM systems of equations.

